# Age- and sex-related topological organisation of human brain functional networks and their relationship to cognition

**DOI:** 10.1101/2021.07.25.453293

**Authors:** Heidi Foo, Anbupalam Thalamuthu, Jiyang Jiang, Forrest Koch, Karen A. Mather, Wei Wen, Perminder S. Sachdev

## Abstract

**BACKGROUND:** Age and sex associated with changes in functional brain network topology and cognition in large population of older adults have been poorly understood. We explored this question further by examining differences in 11 resting-state graph theory measures with respect to age, sex, and their relationships with cognitive performance in 17,127 UK Biobank participants (mean=62.83±7.41 years).

**METHODS:** Brain connectivity toolbox was used to derive the graph theory measures that assessed network integration, segregation, and strength. Multiple linear regression was performed the relationship between age, sex, cognition, and network measures. Subsequently, multivariate analysis was done to further examine the joint effect of the network measures on cognitive functions.

**RESULTS:** Age was associated with an overall decrease in the effectiveness of network communication (i.e. integration) and loss of functional specialisation (i.e. segregation) of specific brain regions. Sex differences were also observed, with women showing more efficient networks which were less segregated than in men (FDR adjusted *p*<.05). Age-related changes were also more apparent in men than women, which suggests that men may be more vulnerable to cognitive decline with age. Interestingly, while network segregation and strength of limbic network were only nominally associated with cognitive performance, the network measures collectively were significantly associated with cognition (FDR adjusted *p*≤.002). This may imply that individual measures may be inadequate to capture much of the variance in neural activity or its output and need further refinement.

**CONCLUSION:** The complexity of the functional brain organisation may be shaped by an individual’s age and sex, which ultimately may influence cognitive performance of older adults. Age and sex stratification may be used to inform clinical neuroscience research to identify older adults at risk of cognitive dysfunction.

## INTRODUCTION

The brain is topographically organised into distinct networks. In the recent years, neuroscientists have examined networks to understand brain function in preference to the classic study of specific brain regions. There are several approaches to mapping these brain networks, with one approach being resting-state functional magnetic resonance imaging (rs-fMRI). Rs-fMRI measures spontaneous brain activity as low-frequency fluctuations in bold oxygen level-dependent (BOLD) signals and is used to understand brain function (1). In network models of rs-fMRI data, functional brain networks are summarised into a collection of nodes (i.e., brain regions) and edges (i.e., magnitude of temporal correlation in fMRI activity between regions) (2, 3). This network model can then be used to study the global and local properties of the functional brain networks (Table 1). There is evidence that adult human brains are organised into groups of specialised functional networks that are able to respond to various cognitive demands (1). Therefore, studying the organisation of functional networks in the ageing brain may allow us to understand age-associated cognitive changes, even in the absence of brain disease (4, 5).

**Table 1.** Graph theory measures and their association with age and age-related diseases

| **Graph theory measures** | **Definition** | **Associations with age and ageing-related diseases** |
| --- | --- | --- |
| Global efficiency | How effectively the information is transmitted at a global level and is the average inverse shortest path length. Higher values imply greater efficiency. | Older age was associated with reduced global efficiency compared to younger participants (12) |
| Characteristic path length | It is the average of all the distances between every pair of nodes in the network. It reflects the integrity of the network and how fast and easily information can flow within the network. Shorter characteristic path length reflects more efficient transmission of information. | Older age was associated with longer characteristic path lengths compared to younger participants (13) |
| Louvain Modularity | Community detection method, which iteratively transforms the network into a set of communities or modules, each consisting of a group of nodes. Higher modularity values indicate denser within-modular connections but sparser connections between nodes that are in different modules. | Brain networks in the elderly showed decreased modularity (less distinct functional networks) but findings were mixed (10) |
| Transitivity | Total of all the clustering coefficients around each node in the network and is normalized collectively. Higher values represent greater specialisation of the brain. | Patients with Alzheimer's disease (AD) showed lower normalized clustering coefficient (i.e. transitivity) (62) |
| Strength | Sum of all neighbouring edge weights. High connectivity strength indicates stronger connectivity between the regions. | Age-related differences were observed in network-level functional connectivity such as increases in auditory network, decreases in connectivity in the visual, frontoparietal, dorsal attention, and salience network. However, findings were mixed (6, 7, 9). |

Reorganisation of the functional networks in the brain has been observed with ageing, and is also associated with changes in cognition (6–10). Age-related alterations have been associated with a less efficient global network, decreased modularity, longer path lengths, and higher clustering coefficient, which may suggest a shift to more local organisation in older age (6, 8, 11, 12). These topological functional network changes occurred most pronouncedly in regions important for cognition. For instance, high clustering coefficients in some frontal, temporal, and parietal regions were related to lower performance in verbal and visual memory functions (13). Declines in default mode network, which comprises the medial and lateral parietal, medial prefrontal, and medial and lateral temporal cortices (14), are reported in ageing and have been associated with memory consolidation (15). In addition, it has been observed that age has a mediating role in the correlation between local clustering coefficients and verbal memory learning scores (13). Similarly, another study found that the relationship between aging and general decline in cognition could be mediated by changes in the functional connectivity measures such as path length (16).

Previous studies have also shown sex differences in the organisation of brain functional networks using graph theory measures. Men showed network segregation (i.e., specialised processing of the brain at a local level) whereas women showed more network integration (i.e., how rapidly the brain can integrate specialised information at a global network level) (8). Another study observed that men had a higher clustering coefficient in the right hemisphere than the left hemisphere (17), suggesting that men had greater specialisation of the right hemisphere. In addition, age-related differences in reorganisation of functional connectivity may also differ by sex, with men showing increasing between-network connectivity (18) while women exhibiting less age-related decreases in the default mode and limbic networks (19). It is noteworthy that age-related changes in cognition also differ by sex. For instance, a recent study has observed that while women had significantly higher baseline global memory, executive function, and memory performance than men, they showed significantly faster declines in the global memory and executive function (20). Another study found that older men had steeper rates of decline on measures of perceptuomotor speed and integration as well as visuospatial abilities (21). Taken together, the findings show that sex may influence age-related functional reorganisation in the brain and improving our understanding of this may shed light onto why some cognitive abilities differ substantially by sex (22).

There is evidence to show that changes in cognition may be due to the changes in functional network connectivity. Segregated functional networks, for instance, seemed to be associated with better long-term episodic memory and fluid processing (23). However, there have been mixed findings regarding how resting-state functional connectivity differences relate to cognitive performance. One longitudinal study found age-related decline of within-network connectivity in default mode and executive control networks but without associations with cognitive decline, whereas an association of between-network connectivity of default mode network and executive control network with processing speed was also observed (24). In contrast, another longitudinal study showed positive associations between within-network connectivity of the default mode network and memory performance (25).

One previous study has investigated the functional network architecture of older adults with respect to age, sex, and cognitive performance (attention, episodic and working memory, executive function, and language) in a cohort of 722 participants with ages between 55 and 85 years old (mean age of 67.1 years) (26). They found resting-state functional connectivity reorganisation with age, particularly in the visual and sensorimotor networks, which may suggest that these networks may mediate age-related differences in cognitive performance. In addition, the authors observed that men showed higher network integration whereas women showed more segregation, which may possibly facilitate sex-related differences in cognitive performance.

This study aims to extend previous work by firstly examining age, sex, and cognitive function in association with functional network properties but in a much larger sample of 17,127 UK Biobank participants. Additionally, a more extensive range of graph theory measures, which assess the global and local properties as well as the strength of the network, will be examined as summarised in Table 1. These measures are namely global efficiency, characteristic path length, Louvain modularity, transitivity, and strength of default, dorsal attention, frontoparietal, limbic, salience, somatomotor, and visual networks, which are typically found to change with aging (7) and are involved in multiple neuropathological processes (27–29).

## METHODS AND MATERIALS

### Participants

Data from 20,598 participants of European ancestry with rs-fMRI scans from the UK Biobank (aged between 44 and 80 years old) (30) were accessed in March 2019. The imaging assessment took place at three different assessment centres: Manchester, Newcastle, and Reading, UK. This project was approved by the NHS National Research Ethics Service (approval letter dated 17th June 2011, ref. 11/NW/0382), project 10279. All data and materials are available via UK Biobank (http://www.ukbiobank.ac.uk).

### Image pre-processing and graph theory analyses

All participants underwent rs-fMRI scan on a Siemens Skyra 3T scanner (Siemens Medical Solutions, Erlangen, Germany). Rs-fMRI was obtained using a blood-oxygen level dependent (BOLD) sequence using am echo-planar imaging (EPI) sequence (TR = 0.735s, TE = 39 ms, FoV = 88 × 88 × 64, voxel resolution 2.4 × 2.4 × 2.4 mm), lasting for ~6 mins (for more details, see https://biobank.ctsu.ox.ac.uk/crystal/crystal/docs/brain_mri.pdf). We analysed the rs-fMRI data that were previously pre-processed by the UK Biobank (31). The pre-processing steps involved: motion correction, intensity normalisation, high-pass temporal filtering, EPI unwarping, and gradient distortion correction. ICA+FIX processing (32–34) was then used to remove structural artefacts. Participants with motion of > 2mm/degrees of translation/rotation were removed. After image pre-processing and quality control, 18,500 participants remained.

The regions of interest (ROIs) used to construct the network properties were selected from the Schaefer atlas (35) corresponding to 100 cortical regions classified into seven resting-state networks including frontoparietal control (FPCN), default mode (DMN), dorsal attention (DAN), salience ventral attention (SVAN), limbic (LIMB), somatomotor (SM), and visual (VIS) networks. 3dNetCorr command from Analysis of Functional Neuroimaging (AFNI) (36) was used to produce network adjacency matrix for each participant. The mean time-series for each region was correlated with the mean time-series for all other regions and extracted for each participant. Further, these time courses are used to estimate the size of signal fluctuation in each node, as well as to estimate connectivity between pairs of nodes using L2-regularisation (rho = 0.5 for Ridge Regression option in FSLNets). More details can be found in Miller, Alfaro-Almagro (37). Partial correlation, *r*, between all pairs of signals was computed to form a 100-by-100 (Schaefer atlas) connectivity matrix, which was then Fisher z-transformed. Self-connections and negative correlations were set to zero. As rs-fMRI can vary across magnitude, the use of undirected weighted matrices may provide a more comprehensive picture of the functional brain networks. The stronger the weights, the stronger the connections between nodes. In addition, we used undirected graph because rs-fMRI data do not permit inferences about the possible direction of information flow. However, undirected graph is useful as it allows us to identify existing connections between specific pairs of network nodes (38). Therefore, we used weighted undirected matrices in our study.

All graph theory measures were derived using the Brain Connectivity Toolbox (BCT) (2). Functional integration can be assessed by global efficiency, which refers to the transmission of information at a global level, and characteristic path length, which is the average shortest distance between any two nodes in the network. To assess network segregation, which characterizes the specialized processing of the brain at a local level, we calculated the Louvain modularity and transitivity. Louvain modularity is a community detection method, which iteratively transforms the network into a set of communities, each consisting of a group of nodes. Higher modularity values indicate denser within-modular connections but sparer connections between nodes that are in different modules. Transitivity refers to the sum of all the clustering coefficients around each node in the network and is normalized collectively. Finally, strength (weighted degree) is described as the sum of all neighbouring edge weights. High connectivity strength indicates stronger connectivity between the regions, which provides an estimation of functional importance of each network. Subsequently, we averaged the two hemispheres to derive a value for each node and averaged within each network to derive a value for each of the 7 networks for strength measures. A total of eleven graph theory measures were used in the current study.

### Cognition

Cognitive assessments were administered on a touchscreen computer and were acquired at the imaging visit (instance 2). Seven tests from the UK Biobank battery of tests were selected to represent three cognitive domains(39, 40) namely processing speed, memory, and executive function, in this study. All test scores were first z-transformed and then averaged to form domain scores. Processing speed domain included the following tests: “Reaction Time” (average time to correctly identify matches in a “snap”-like card game task), “Trail Making A” (time taken to complete a numeric path), and “Symbol Digit Substitution” (number of correct symbol number matches within the time limit). “Numeric Memory” (maximum number of digits remembered correctly) and “Pairs Matching” (number of incorrect visual matching) represented the memory domain whereas “Trail Making B” (time taken to complete an alphanumeric path) and “Fluid Intelligence” (total number of questions that required logic and reasoning correctly answered) formed the executive function domain. Global cognition was computed by averaging the domain scores and z-transformed. After including those with cognition and graph theory data, the final sample in this study was 17,127 UK Biobank participants.

### Statistical analysis

Statistical analyses were performed with R (V 4.0.0) (41). The graph theory measures were normalized using ranked transformed within the ‘rntransform’ function in R from GeneABEL package (42) and age was z-transformed for regression analysis. In line with previous studies (43), we controlled for imaging covariates, including head size, head motion from rs-fMRI, and volumetric scaling factor needed to normalize for head size, as well as scanning site and education. The network measures were residualised for imaging covariates and assessment centre and used in all subsequent analyses.

To explore age effects and sex-related changes in the networks, a multiple linear regression that modelled the targeted property of networks as the dependent variable and age, age^2^, sex (Female = 0, Male = 1), years of education, and age-by-sex and age^2^-by-sex interactions as predictors was undertaken. In addition, separate multiple linear regressions were performed to study whether the network measures influenced cognitive functions (dependent variable) with covariates as in the previous model.

Multivariate analysis was done to further examine the joint effect of the network measures on cognitive functions after accounting for the same set of covariates in the univariate model. Since the network measures are correlated we used penalised regression analysis using glmnet algorithm as implemented in the r package caret (44). The glmnet uses two penalty functions with tuning parameters to shrink the beta coefficients in the generalised linear model (glm). We used elastic net glm model with default options to identify the optimum tuning parameter estimates. Network measures and the covariates with non-zero regression co-efficient in the training step was fit with linear regression model. Likelihood ratio tests, p-values and the incremental r-square were computed by comparing the model with network measures (full model) again a model with only the covariates (base model). False discovery rate – adjusted p-values were obtained by using Benjamini and Hochberg (45) procedure as implemented in the R function *p.adjust.*

## RESULTS

### Sample characteristics

The current sample of 17,127 participants is a group of generally healthy middle-aged and older adults (range = 45.17 – 80.67 years, mean age = 62.83 ± 7.41 years) after including only samples with cognition and graph theory data. Of this sample 9,037 were women and 8,090 were men, with an overall mean of 15.73 (± 4.74) years of education. Significant differences were observed for the demographics, graph theory measures, and memory scores between men and women (Table 2). Figure 1 shows the significant correlations between the network measures, except for transitivity, which was not significantly associated with any other measures.

**Table 2.**
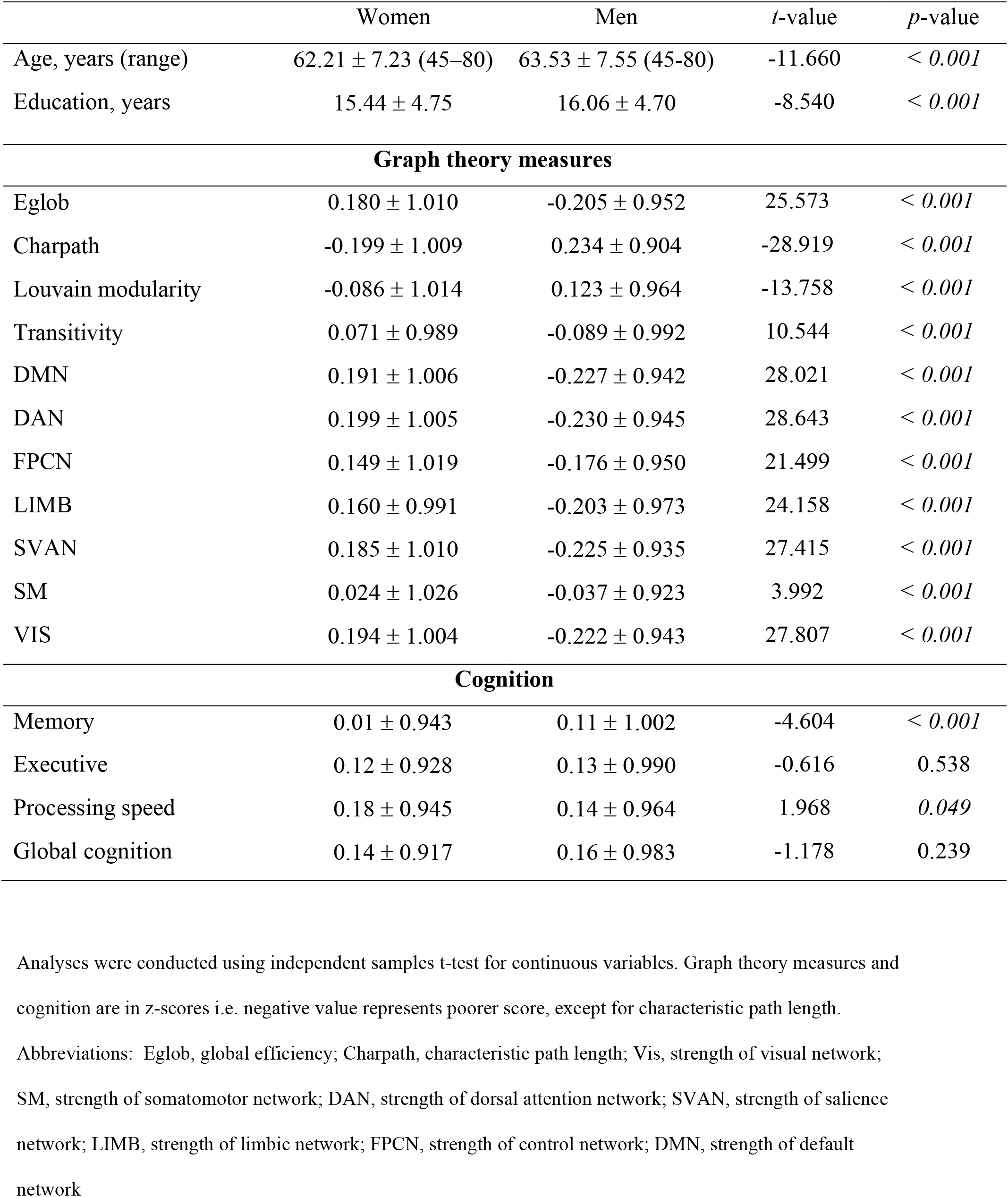
UK biobank sample characteristics and descriptive statistics (mean ± standard deviation) of graph theory measures and cognition measures in women and men

|  | Women | Men | *t*-value | *p*-value |
| --- | --- | --- | --- | --- |
| Age, years (range) | 62.21 ± 7.23 (45–80) | 63.53 ± 7.55 (45-80) | -11.660 | < 0.001 |
| Education, years | 15.44 ± 4.75 | 16.06 ± 4.70 | -8.540 | < 0.001 |
| **Graph theory measures** |  |  |  |  |
| Eglob | 0.180 ± 1.010 | -0.205 ± 0.952 | 25.573 | < 0.001 |
| Charpath | -0.199 ± 1.009 | 0.234 ± 0.904 | -28.919 | < 0.001 |
| Louvain modularity | -0.086 ± 1.014 | 0.123 ± 0.964 | -13.758 | < 0.001 |
| Transitivity | 0.071 ± 0.989 | -0.089 ± 0.992 | 10.544 | < 0.001 |
| DMN | 0.191 ± 1.006 | -0.227 ± 0.942 | 28.021 | < 0.001 |
| DAN | 0.199 ± 1.005 | -0.230 ± 0.945 | 28.643 | < 0.001 |
| FPCN | 0.149 ± 1.019 | -0.176 ± 0.950 | 21.499 | < 0.001 |
| LIMB | 0.160 ± 0.991 | -0.203 ± 0.973 | 24.158 | < 0.001 |
| SVAN | 0.185 ± 1.010 | -0.225 ± 0.935 | 27.415 | < 0.001 |
| SM | 0.024 ± 1.026 | -0.037 ± 0.923 | 3.992 | < 0.001 |
| VIS | 0.194 ± 1.004 | -0.222 ± 0.943 | 27.807 | < 0.001 |
| **Cognition** |  |  |  |  |
| Memory | 0.01 ± 0.943 | 0.11 ± 1.002 | -4.604 | < 0.001 |
| Executive | 0.12 ± 0.928 | 0.13 ± 0.990 | -0.616 | 0.538 |
| Processing speed | 0.18 ± 0.945 | 0.14 ± 0.964 | 1.968 | 0.049 |
| Global cognition | 0.14 ± 0.917 | 0.16 ± 0.983 | -1.178 | 0.239 |

Analyses were conducted using independent samples t-test for continuous variables. Graph theory measures and cognition are in z-scores i.e. negative value represents poorer score, except for characteristic path length.

**Figure 1.**
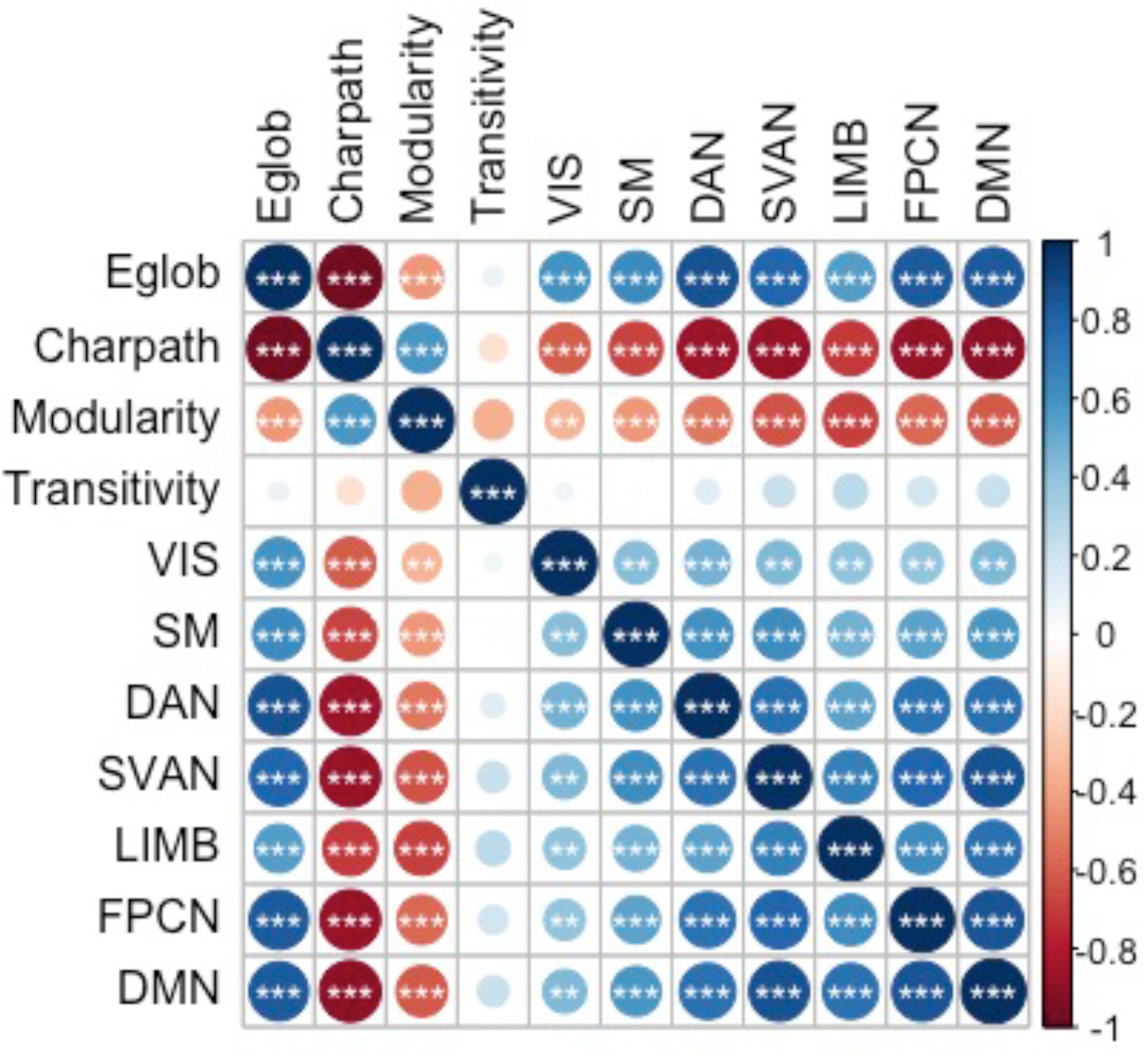
Correlations between the graph theory measures * *p* < 0.05, ** *p* < 0.01, *** *p* < 0.001 Abbreviations: Eglob, global efficiency; Charpath, characteristic path length; Vis, strength of visual network; SM, strength of somatomotor network; DAN, strength of dorsal attention network; SVAN, strength of salience network; LIMB, strength of limbic network; FPCN, strength of control network; DMN, strength of default network

### Age- and sex- related differences in functional brain network

Figure 2 and Table 3 summarise the results of age- and sex- related differences in the graph theory measures. Global efficiency, Louvain modularity, and strength of the networks decreased significantly with age, whereas characteristic path length and transitivity increased significantly with age. The only exceptions were that strength of default and salience networks were not significantly associated with age.

**Figure 2.**
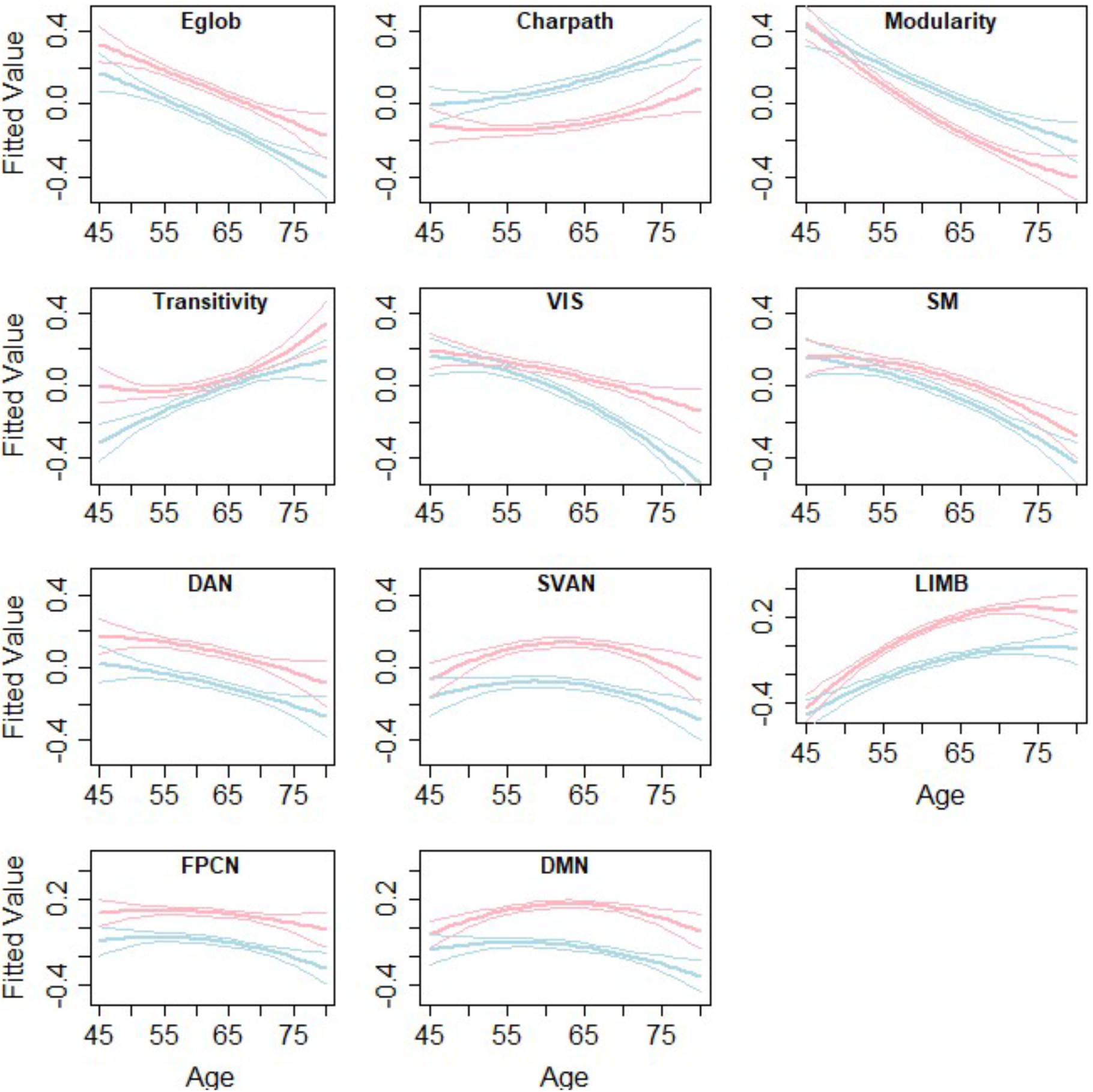
Age- and sex-related differences in the graph theory measures Lines represent the fitted values for men (blue) and women (red) separately. The middle line shows the fitted equation evaluated at the mean value of education for each sex, while the top and lower lines represent confidence bands. Abbreviations: Eglob, global efficiency; Charpath, characteristic path length; Vis, strength of visual network; SM, strength of somatomotor network; DAN, strength of dorsal attention network; SVAN, strength of salience network; LIMB, strength of limbic network; FPCN, strength of control network; DMN, strength of default network

**Table 3.** Age- and sex-related differences in graph theory measures

| Graph theory measures | $\beta$<br>Age | SE<br>Age | $\beta$<br>Sex | SE<br>Sex | $\beta$<br>AgeXsex | SE<br>AgeXsex | *Padj*<br>Age | *Padj*<br>Sex | *Padj*<br>AgeXsex |
| --- | --- | --- | --- | --- | --- | --- | --- | --- | --- |
| Eglob | -0.108 | 0.011 | -0.170 | 0.021 | -0.014 | 0.016 | 3.74E-21* | 1.29E-15* | 0.391 |
| Charpath | 0.043 | 0.011 | 0.226 | 0.021 | 0.034 | 0.016 | 1.67E-04* | 3.78E-26* | 0.068 |
| Louvain modularity | -0.181 | 0.011 | 0.166 | 0.021 | 0.046 | 0.016 | 1.05E-57* | 2.96E-15* | 0.016* |
| Transitivity | 0.072 | 0.011 | -0.042 | 0.021 | 0.024 | 0.016 | 4.55E-10* | 0.048* | 0.179 |
| DMN | 0.004 | 0.011 | -0.287 | 0.021 | -0.044 | 0.016 | 0.772 | 4.28E-41* | 0.016* |
| DAN | -0.055 | 0.011 | -0.181 | 0.021 | -0.006 | 0.016 | 1.71E-06* | 2.27E-17* | 0.681 |
| FPCN | -0.025 | 0.011 | -0.194 | 0.021 | -0.015 | 0.016 | 0.032* | 1.11E-19* | 0.391 |
| LIMB | 0.142 | 0.011 | -0.263 | 0.021 | -0.043 | 0.016 | 3.14E-36* | 3.04E-35* | 0.016* |
| SVAN | 0.000 | 0.011 | -0.223 | 0.021 | -0.026 | 0.016 | 0.971 | 2.29E-25* | 0.145 |
| SM | -0.093 | 0.011 | -0.090 | 0.021 | -0.031 | 0.016 | 5.86E-16* | 2.27E-05* | 0.093 |
| VIS | -0.071 | 0.011 | -0.106 | 0.021 | -0.076 | 0.016 | 5.12E-10* | 6.80E-07* | 1.21E-05* |

\* represents significance
Abbreviations: $\beta$ = beta; SE, standard error, *Padj*, adjusted *p*-value; AgeXsex, age and sex interaction; Eglob, global efficiency; Charpath, characteristic path length; Vis, strength of visual network; SM, strength of somatomotor network; DAN, strength of dorsal attention network; SVAN, strength of salience network; LIMB, strength of limbic network; FPCN, strength of control network; DMN, strength of default network

Sex was significantly associated with all measures, except for transitivity. Men appeared to have lower global efficiency, transitivity, and strengths of all the networks as well as longer characteristic path lengths compared to women. Men showed increased Louvain modularity compared to women.

Age and sex interaction were negatively associated with Louvain modularity, and strength of visual, limbic, and default networks. This implies that age-related changes in these measures were more apparent in males than females.

### Association of network measures with cognition

We examined the network influence on cognition after controlling for age, age^2^, sex, and education. Although none of these results would survive correction for multiple testing, we report the results that were nominally significant. Louvain modularity showed positive associations with global cognition whereas transitivity was negatively associated with memory. Strength of limbic network also showed negative associations with global cognition and memory (Supplementary table 1).

We further examined this relationship to see if it was moderated by age and sex. However, none of the interaction effects between network measures and age or sex on cognition were significant (Supplementary table 2).

### Multivariate analysis between network measures and cognition

Given the significant correlations between the network measures, we further investigated whether the joint effect of the network measures contributed to cognition after controlling for age, sex, and education variables. A summary of the results of each of the examined models are presented in Supplementary table 3. We observed that while the R^2^ difference between the base model (age, age^2^, sex, and education) and the full model with the network measures was small, the joint effect of the network measures still significantly contributed to cognition (Table 4).

**Table 4.** Multivariate analysis of the joint effect of the network measures with cognitive function

| **Cognitive domains** | **df** | **LR** | **R<sup>2</sup> full model** | **R<sup>2</sup> base** | **R<sup>2</sup> diff** | ***P*adj** |
| --- | --- | --- | --- | --- | --- | --- |
| Processing speed | 7 | 3.224 | 0.239 | 0.237 | 0.002 | *0.002* |
| Executive function | 9 | 3.503 | 0.132 | 0.128 | 0.004 | *5.29E-04* |
| Memory | 8 | 3.650 | 0.045 | 0.041 | 0.004 | *5.29E-04* |
| Global cognition | 3 | 4.480 | 0.188 | 0.184 | 0.004 | *7.75E-05* |

Abbreviations: df, number of network measures in the model; LR, likelihood ratio, diff, difference; *P*adj, adjusted p-value
Networks included in the final model:
Processing speed – Age, Age2, Sex, Education, Louvain Modularity, Transitivity, Strength of Visual Network, Strength of Somatomotor Network, Strength of Dorsal Attention Network, Strength of Salience Network, Strength of Limbic Network
Executive function - Age, Age2, Sex, Education, Louvain Modularity, Transitivity, Strength of Visual Network, Strength of Somatomotor Network, Strength of Dorsal Attention Network, Strength of Salience Network, Strength of Limbic Network, Strength of Control Network, Strength of Default Network
Memory - Age, Age2, Sex, Education, Global efficiency, Transitivity, Strength of Visual Network, Strength of Dorsal Attention Network, Strength of Salience Network, Strength of Limbic Network, Strength of Control Network, Strength of Default Network
Global cognition - Age, Age2, Sex, Education, Louvain Modularity, Transitivity, Strength of Visual Network, Strength of Somatomotor Network, Strength of Dorsal Attention Network, Strength of Limbic Network, Strength of Control Network, Strength of Default Network
Full model includes network measures and base model includes only covariates

## DISCUSSION

Changes to resting-state networks due to ageing arguably reflect more fundamental alterations or adaptations at the general level of brain function (46). Graph theoretical approaches may be the most integrative way to investigate resting-state functional connectivity (RSFC) as it studies connectivity at both nodal and systems levels (46). Therefore, in this study, we examined the topological age and sex relationship with functional brain networks, using graph theory measures, and cognition. We observed that most functional brain network measures showed decreasing strength of connectivity as well as reduced efficiency of communication and specialisation between the networks with ageing. However, the default mode and salience networks were an exception to this finding, with no significant results observed. In addition, there were significant sex differences in brain functional network topology where women showed greater efficiency of networks and network strength but less modularity than men. Further, age-related changes were more apparent in men than women. Lastly, the collective effect of the network measures contributed significantly to cognitive performance, with the highest correlation being with processing speed. However, no one network measure was significant after multiple testing adjustment.

We observed that global efficiency correlated negatively with age whereas characteristic path length correlated positively with age, which was similar to a previous study (13). This suggests an overall age-related decrease in the effectiveness of the communication between brain regions. In addition, the finding that modularity decreases with age has also been reported previously (7). This implies that increasing age is associated with a less differentiated functional modular structure, which may be either due to the increase in between-network connections or the decrease in within-network connections or both (7, 10). At younger ages, functional brain networks are more segregated with every network being relatively specialised for distinct mental processes (10). The data suggest that there is some loss of functional specialisation of specific brain networks as the brain ages (47), which may be important for cognitive reserve and compensation in older adults. . Furthermore, we showed age-related decline in all of the other network strengths, excluding the DMN and salience network. However, results for other networks from previous studies are more complex. For instance, Betzel and colleagues (9) found within-network decline for higher order control and attention networks but stability for visual and somatomotor networks, whilst another study (7) showed increased global and local efficiency in the sensorimotor network in older compared to younger adults. Taken together, our data and others suggest age-related vulnerability in global network measures as well as specific network strengths.

Importantly, we did not observe any age-related decline in the DMN and salience network. Prior works suggest that within-network posterior DMN connectivity, including the angular gyrus, anterior cingulate cortex, precuneus, dorsal prefrontal, and inferior parietal lobe, decreases with age, (6, 7, 9, 10, 26). In contrast, within the older adult population, DMN as a whole remains relatively stable (26, 48). This finding is important as it shows that anterior-posterior DMN has differential vulnerability to age-related changes. Moreover, the salience network seems to remain relatively stable throughout the lifespan (10, 49) as well as in older age (26, 50). Interestingly, the DMN and salience network have also been implicated in age-related diseases such as Alzheimer’s disease (AD) and depression. One study observed that individuals with AD showed moderate decrease of within-network DMN between the posterior cingulate cortex and right hippocampus as compared to healthy controls but no differences were evident for whole-network DMN (51). Further, compared to older adult controls, individuals with AD showed significantly decreased within-network functional connectivity in the frontoinsular cortices and increased FC in medial prefrontal cortex in the salience network (52). Similarly, older adults with depression demonstrated higher within-network DMN in the left precuneus, subgenual anterior cingulate cortex (ACC), ventromedial prefrontal cortex, and lateral parietal regions than controls (53). In addition, regarding the salience network, within-network bilateral anterior insula showed decreased connectivity but bilateral ACC showed increased connectivity in middle-aged adults with depression compared to controls (54). These findings suggest that while the whole DMN may be preserved, within-network posterior DMN may be vulnerable to ageing and ageing-related diseases.

The topology of functional brain networks differed by sex. We detected significant sex effects on all the assessed graph theory measures. Consistent with results from Zhang et al. (8) showing that female brains facilitated functional integration in young adults, we found that in older individuals, women indeed had higher global efficiency and shorter characteristic path length than men. Similarly, congruent with previous findings, we also observed women had higher normalised clustering coefficients (i.e. transitivity) than men (8). However, men exhibited stronger Louvain modularity which suggests that there may be sex differences even within network segregation. It has previously been reported that women tend to exhibit overall higher within-network RSFC (55), which is consistent with our finding that women had higher network strengths than men. Similarly, consistent with previous findings that women show less age-related decreases in RSFC in the default and limbic network (19), we found that age-related changes in strengths of the limbic and default networks in addition to Louvain modularity and strength of visual network were more apparent in males than females. This suggests that ageing-related changes in the functional brain network are different in the two sexes and that this difference may in part account for the differential vulnerability in cognitive decline between men and women.

Functional connectivity architecture in the brain has been associated with cognitive performance in older adults independent of age, sex, and education in this study. We observed that decreased Louvain modularity was nominally associated with decline in global cognition and that decreased transitivity was nominally associated with decline in memory. Individuals with less segregated networks exhibited poorest memory ability after controlling for age, which may suggest that network segregation may be an age-invariant marker of individual differences in cognition (10). In addition, prior evidence from cognitive training interventions has shown that higher modularity at baseline in older adults was associated with greater cognitive training improvements, especially in sensory-motor processing (56). Furthermore, given that the limbic network derived from the Schaefer parcellation comprises the orbitofrontal cortex and temporal pole, and these regions are associated with memory formation (57) and executive function (58), it supports our finding that the strength of the limbic network showed negative associations with memory and executive function. While there is nominal significance between individual network measures and cognition, the joint effect of all the network measures contributed significantly to cognition after accounting for age, sex, and education. This suggests that cognitive decline observed in older adults may be partially explained by independent changes in brain functional network organisation. It also implies that individual network measures may be inadequate to capture much of the variance in neural activity and the functional output. Future studies are needed to combine various strategies to more holistically understand the network topology in relation to cognition.

The strengths of this study include a well-characterised large middle and older aged cohort, uniform imaging methods, the inclusion of a range of network measures associated with age and ageing-related diseases, and the examination of a number of cognitive domains. This is the largest study of its kind thus far. However, limitations should also be considered. Firstly, this study is cross-sectional, which precludes the ability to detect subtle changes in the functional brain topology over time within individuals. Secondly, while using weighted undirected matrix circumvents issues surrounding filtering/thresholding the connectivity matrix to maintain significant edge weights represented in a binary matrix, there are inherent difficulties associated with the interpretation of the results. As brain signals recorded from resting-state fMRI are typically noisy, it is possible that edge weights may be affected by non-neural contributions (59). Despite this, with careful denoising of the resting-state fMRI data (60, 61) and covarying for motion, it is possible to minimize the noise in the data. Given that we have performed motion correction and included it as a covariate as well as performed regularisation on the imaging data, we are confident that the estimation of the partial correlation matrix derived for subsequent analysis of the graph theory measures is valid. Moreover, while we were only interested in investigating whole network functional connectivity, given the findings from DMN and salience network, it may be beneficial to look at individual nodes within the network to more holistically capture the nodal topology. Lastly, given the principles of neurobiology, we assumed that network properties influence cognition and not the other way around. This question needs to be examined longitudinally to confirm the directionality of the relationship.

In conclusion, in this large population-based study age was associated with decreased overall network integrity and specialised processing of the brain at a local level. Women had better functional network topology properties than men, with men tending to have denser within-network connections but sparser connections between-network connections. This work demonstrates the complexity of functional brain organisation that is shaped by age, sex, and other factors, which ultimately may influence cognitive performance of older adults.

## ACKNOWLEDGMENTS AND DISCLOSURES

This research did not receive any specific grant from funding agencies in the public, commercial, or not-for-profit sectors. Thanks to the UNSW Scientia Scholarship Programme for their support to HF. The authors report no biomedical financial interests or potential conflicts of interest.

